# Redirecting vacuolar nitrate transport improves nitrogen use efficiency and seed protein content

**DOI:** 10.64898/2026.08.21.746244

**Authors:** Anne Marmagne, Yannick Fierlej, Benoit Bernay, Caroline Cukier, Jérémy Lothier, Céline Masclaux-Daubresse, Fabien Chardon

## Abstract

Improving seed protein content without compromising carbon allocation or yield is a major challenge for enhancing nitrogen use efficiency. Here, we show that redirecting vacuolar nitrate transport through concurrent manipulation of tonoplast proteins controlling nitrate storage or export provides an effective lever to reprogram nitrogen allocation from leaves toward the seeds. Using *Arabidopsis thaliana Ws* lines disrupted for the vacuolar CLC-a nitrate importer and/or overexpressing the *NRT2.7* tonoplast nitrate exporter, we show that plants combining the two modifications (*35S::NRT2.7(clc-a))* integrate reduced nitrogen retention in vegetative tissues with increased nitrogen allocation to seeds. As a result, *35S::NRT2.7(clc-a)* plants exhibit the strongest increase in seed protein content among all genotypes (approximately +25%) without affecting seed yield, carbon concentration, or lipid composition. Altered vacuolar nitrate fluxes in *35S::NRT2.7(clc-a)* stimulate nitrate assimilation, enhance nitrate reductase activity and amino acid biosynthetic pathways, and drive coordinated reprogramming of nitrogen and carbon metabolisms. Through ^15^N pulse–chase experiments, we confirmed that *35S::NRT2.7(clc-a)* shows the highest nitrogen remobilization efficiency toward seeds. Overexpression of the barley *NRT2.7* homolog *HvNRT2.10* in Arabidopsis wild type and clc-a backgrounds reproduces the key features of *35S::NRT2.7* phenotype, demonstrating the conservation of *NRT2.7* regulatory effects on plant metabolism across species. Together, these findings identify vacuolar nitrate transport as a promising target to modulate grain protein content in cereals through genetic strategies acting on nitrogen storage and remobilization.

**HIGHLIGHTS:**

- Vacuolar nitrate transport modifications (*clc-a, 35S::NRT2.7*, *35S::NRT2.7(clc-a)*) increase seed nitrogen and protein concentrations without altering seed yield or carbon levels, highlighting a selective enhancement in nitrogen storage.
- *NRT2.7* overexpression (especially in the *clc-a* background) redirects nitrogen resources from vegetative tissues to the seeds, emphasizing the role of NRT2.7 in nitrogen remobilization.
- Proteomic analyses revealed distinct and contrasting profiles between *clc-a* and *35S::NRT2.7*, with *35S::NRT2.7(clc-a)*, exhibiting unique effects in auxin transport and immunity pathways.
- Overexpression of *barley homolog of AtNRT2.7, HvNRT2.10*, in Arabidopsis confirmed conserved role for NRT2.7 family in boosting seed nitrogen concentration and NUE in wild type and in a larger extent in the *clc-a* background.

## INTRODUCTION

Nitrate is the primary form of nitrogen (N) acquired from the soil by most plant species (Wang et al., 2012). As an essential macronutrient, nitrogen supports plant growth and seed production and can be either stored in vacuoles or assimilated into amino acids and more complex biomolecules. In annual plants such as Arabidopsis (*Arabidopsis thaliana* L.), approximately 40% of seed nitrogen originates from post-flowering nitrate uptake, while the remaining portion is remobilized from senescing leaves through nitrogen recycling processes that support seed filling (Marmagne et al., 2022). Because excessive nitrogen fertilization leads to environmental pollution and high production costs (Fowler et al., 2013; Verhoeven et al., 2017), improving nitrogen use efficiency (NUE), defined as the plant’s ability to optimize nitrogen uptake, assimilation, and remobilization, has become a major goal in sustainable agriculture. While significant efforts have focused on enhancing nitrogen uptake and assimilation, the mechanisms controlling nitrogen storage and remobilization remain relatively underexplored (Havé et al., 2017). Gaining a deeper understanding of these processes is crucial for improving NUE and reducing fertilizer dependency in crops.

Despite two decades of research, improving NUE in crops through biotechnological approaches remains a major challenge (Lebedev et al., 2021). NUE is a complex trait governed by a network of genetic and physiological factors, including nitrogen uptake, assimilation, remobilization, and its coordination with carbon metabolism (Marmagne et al., 2022). Various strategies have targeted genes encoding nitrate and ammonium transporters, key enzymes in nitrogen assimilation such as nitrate reductase and glutamine synthetase, or transcriptional regulators. However, many of these approaches have led to unintended side effects, including growth penalties and altered development (The et al., 2021; Lebedev et al., 2021). Although research has largely focused on increasing nitrogen acquisition, these efforts have had limited success when translated to field conditions. Several hypotheses have been proposed to explain these inconsistent outcomes, including negative pleiotropic effects from constitutive promoter usage, imbalances in nitrogen metabolite pools leading to feedback inhibition, substrate limitations for amino acid biosynthesis, or post-transcriptional and post-translational regulation (The et al., 2021). These limitations underscore the need for alternative strategies targeting nitrogen remobilization and allocation, rather than solely uptake and assimilation. In this context, vacuolar nitrate transporters represent a promising but underexplored target for enhancing NUE.

Vacuoles serve as the primary nitrate storage compartment in plant cells, playing a central role in nitrogen homeostasis by buffering fluctuations in external nitrogen availability and regulating intracellular nitrate distribution (Angeli et al., 2008). In Arabidopsis, over 90% of cellular nitrate is sequestered in vacuoles, where it also contributes to osmotic regulation and charge balance (Lu et al., 2022). Maintaining a dynamic equilibrium between vacuolar nitrate accumulation and cytosolic release is essential for coordinating nitrate assimilation and redistributing nitrogen between vegetative and reproductive organs.

Among vacuolar nitrate transporters, the first to be characterized was CLC-a (Chloride Channel-a), a 2NO₃⁻/H⁺ exchanger responsible for nitrate sequestration into vacuoles, especially in leaves (Hechenberger et al., 1996; De Angeli et al., 2006). Loss-of-function *clc-a* mutants exhibit a 50% reduction in vacuolar nitrate content, accompanied by increased activity of nitrate assimilation enzymes such as nitrate reductase and glutamine synthetase, highlighting the essential role of CLC-a in nitrate storage and nitrogen homeostasis (Geelen et al., 2000; Liao et al., 2018). Recent work by Hodin et al. (2023) further demonstrated that CLC-a influences cytosolic nitrate availability and plant growth regulation. During reproductive development, *clc-a* mutants showed altered nitrogen allocation compared to wild-type plants, retaining less nitrogen in rosettes while exhibiting higher nitrogen concentrations in seeds. This shift was associated with enhanced post-flowering nitrate uptake and ultimately led to improved NUE, underscoring the potential of targeting vacuolar nitrate transport to optimize nitrogen distribution in plants. However, in addition to its role in nitrogen metabolism, CLC-a is critical for stomatal function. By contributing to vacuolar nitrate pools, it influences turgor pressure and stomatal closure. Disruption of CLC-a impairs stomatal regulation and reduces plant water content and plant growth, resulting in lower water use efficiency (WUE) (De Angeli et al., 2006; Wege et al., 2014; Hodin et al., 2023). CLC-a belongs to the broader *CLC* gene family, which in Arabidopsis comprises six other members. Among them, AtCLCb, AtCLCc, and AtCLCg are also localized to the tonoplast (Zifarelli and Pusch, 2010; Von Der Fecht-Bartenbach et al., 2010). Although several studies suggest that these isoforms participate in vacuolar nitrate storage or stomatal movement (Harada et al., 2004; De Angeli et al., 2006; Jossier et al., 2010; Nguyen et al., 2016), their precise roles in nitrate homeostasis remain unclear and appear to be genotype-dependent and generally less prominent than that of CLC-a. Together, these findings indicate that CLC proteins have diverse functions in ion homeostasis, stress adaptation, and nitrogen utilization, with CLC-a emerging as a particularly promising target for improving NUE through modulation of vacuolar nitrate transport, despite its known negative effects on plant growth.

The high-affinity nitrate transporter AtNRT2.7 is the second best-characterized vacuolar nitrate transporter in *Arabidopsis*, primarily known for its role in nitrate accumulation in seeds (Chopin et al., 2007; David et al., 2014). Unlike other NRT2 family members localized at the plasma membrane, NRT2.7 is targeted to the tonoplast and is strongly expressed in mature seeds, with weaker expression in leaves and roots (Orsel et al., 2002; Okamoto et al., 2003). Loss of NRT2.7 function markedly reduces seed nitrate content, confirming its importance during the reproductive stage. Recent work has expanded this view by showing that ectopic overexpression of NRT2.7 promotes nitrate efflux from the vacuole and enhances plant growth under low nitrogen, without restoring the nitrate homeostasis defects of *clc-a* mutants (Armengaud et al., 2025). These findings indicate that CLCa and NRT2.7 fulfil distinct and non-redundant roles in vacuolar nitrate management, with CLC-a primarily enabling storage and NRT2.7 contributing to nitrate mobilization when overexpressed.

In contrast to vacuolar nitrate storage, the mechanisms controlling nitrate efflux from the vacuole remain less well defined and likely involve several transporters. Three tonoplast-localized NPF proteins (NPF5.10, NPF5.14, and NPF8.5) have been identified as contributors to nitrate release into the cytosol in the vascular stele of roots and leaves (He et al., 2017). While single mutants do not exhibit strong phenotypes, the *npf5.10 npf5.14 npf8.5* triple mutant shows enhanced nitrate translocation from root to shoot without affecting root nitrate uptake, suggesting that vacuolar nitrate export relies on the coordinated action of multiple transporters. Together with the dual activities of CLC-a and the nitrate-mobilizing potential of NRT2.7, these results highlight the functional diversity of vacuolar nitrate transport systems.

Despite these advances, vacuolar nitrate transport remains a largely underexplored lever for improving NUE. Most strategies aimed at enhancing NUE have focused on nitrate uptake or assimilation, yet recent findings clearly demonstrated that modifying nitrate fluxes across the tonoplast can profoundly influence nitrate homeostasis, biomass production, and nitrogen allocation among organs. However, how the combined manipulation of vacuolar nitrate storage (*via* CLC-A) and vacuolar nitrate export (*via* NRT2.7) influences nitrogen remobilization during reproductive development, seed composition, and overall NUE remains unknown. To address this question, we investigated how manipulating vacuolar nitrate storage and export through *CLC-A* disruption and *NRT2.7* overexpression affects nitrate homeostasis, nitrogen remobilization from vegetative leaves to seeds, and seed nitrogen composition in Arabidopsis. By combining physiological, metabolic, and proteomic analyses, we show that coordinated modulation of vacuolar nitrate fluxes profoundly reshapes nitrogen allocation, modifies seed protein accumulation, and enhances nitrogen use efficiency.

## RESULTS

### Modifications of vacuolar nitrate transport affects nitrogen composition in seeds without altering seed yield or carbon content

Alterations in vacuolar nitrate transport, achieved through disruption of *CLC-A* and/or overexpression of *NRT2.7* did not significantly impact seed yield (Figure 1A). However, N concentration in seeds was notably higher in mutants and overexpressing lines compared to wild-type. The *clc-a* and *35S::NRT2.7(clc-a)* lines both exhibited a 11% increase in seed N concentration (Figure 1B). Interestingly, despite this increase in N, and the well-established negative correlation between N and C concentrations (Marmagne et al., 2020), no significant change in seed C concentration were observed across the lines (Figure 1C). Further analysis of seed composition revealed a 25% increase in seed protein concentration in the *35S::NRT2.7(clc-a)* line, along with a significant rise in nitrate concentration in both *35S::NRT2.7* and *35S::NRT2.7(clc-a)* seeds (+62% and +91% respectively), compared to the wild-type (Figure 1D). In contrast, lipid concentration and composition remained unchanged between the modified lines and wild-type (Figures 1E, S1). Interestingly, the weight of one seed was also higher in the *35S::NRT2.7(clc-a)* line (Figure 1G), though seeds of clc-a and *35S::NRT2.7* were not heavier than that of WT. Protein composition analysis via SDS-PAGE confirmed the larger content of seed storage proteins, particularly the 12S alpha-and 12S beta-globulins, in the *35S::NRT2.7(clc-a)* seeds compared to the wild-type (Figure 1F). Overall, results indicate that modifying CLC-a and NRT2.7 vacuolar nitrate transporters affect seed nitrogen and protein concentrations without impacting seed yield or carbon concentrations. The *35S::NRT2.7(clc-a)* line is of particular interest as it combines higher seed nitrogen concentrations as in *clc-a* and higher seed nitrate concentration as found in *NRT2.7* overexpressing lines. Notably, marked increase in seed storage proteins distinguishing *35S::NRT2.7(clc-a)* from the single modified lines.

**Figure 1.**
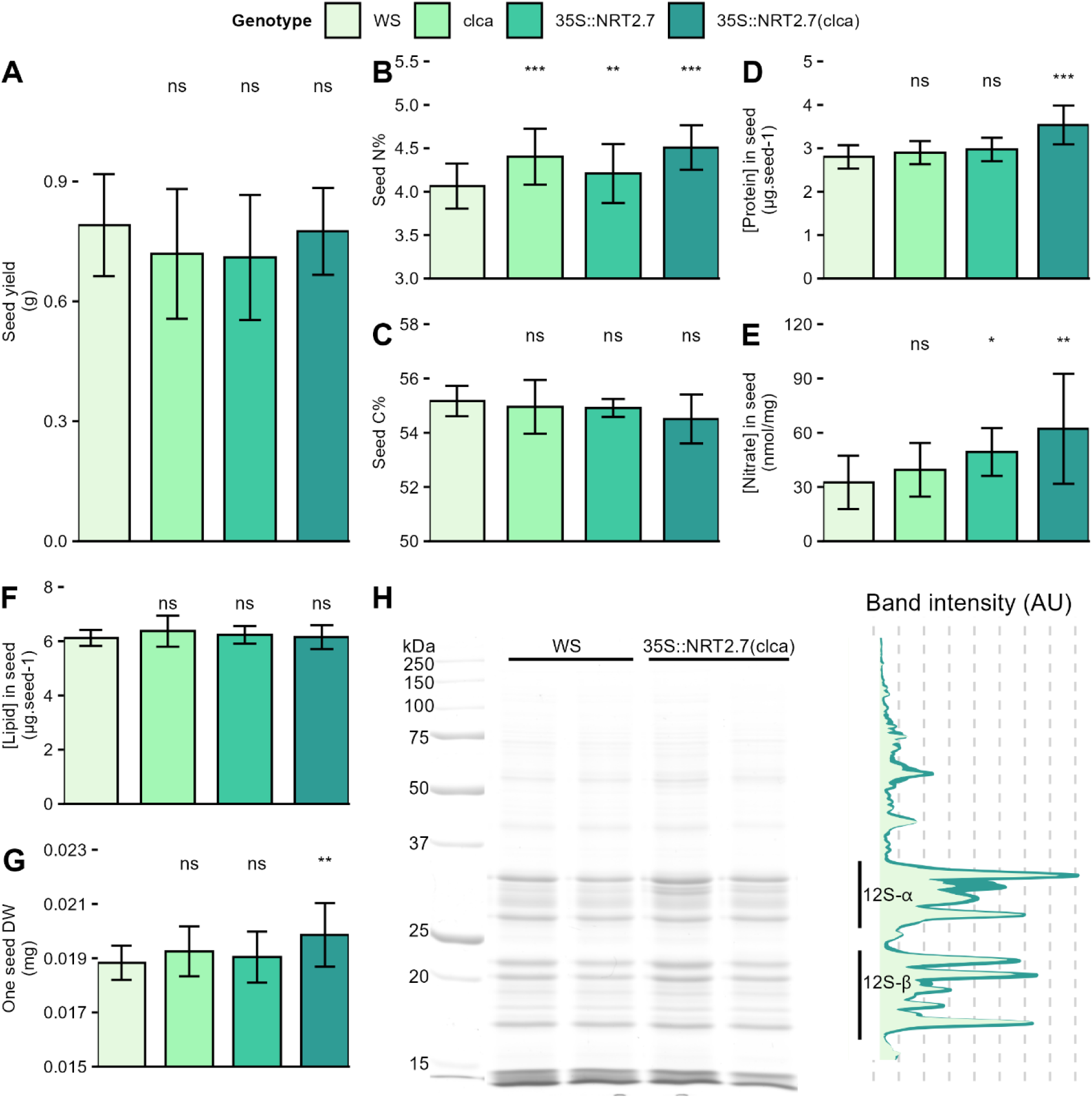
Modification of the vacuolar nitrate transport system impacts nitrogen composition in seeds without disturbing seed yield or carbon concentration in seeds. Plants were grown on sand under high-nitrate condition, in short days for vegetative growth followed by long days for flowering. (A) Seed yield per plant. (B–C) Total nitrogen and carbon concentrations in seeds (N% and C%). (D) Total soluble protein content per seed. (E) Nitrate concentration in seeds. (F) Total lipid content in seeds. (G) Individual seed dry weight. For all the histograms (A-G) bars represent means ± standard deviation, and stars indicate significant differences between WS and the other genotypes (t-test, *p<0.05, **p<0.01, ***p<0.001 n=20). (H) Silver staining of total seed protein extracts after separation on SDS–PAGE.

### Modifications of vacuolar nitrate transport alter nitrogen storage and metabolism in leaves

To better understand how modifications of vacuolar nitrate transport affect leaf metabolism, we harvested the rosettes at vegetative stage, from two independent cultures. Fresh rosette weight was slightly lower in the *clc-a* mutant (1.75 g/plant) compared to the wild type (2.00 g/plant) (Figure 2A). Although differences were not statistically significant, the *35S::NRT2.7* line tended to have heavier rosettes, while rosettes of *35S::NRT2.7(clc-a)* tended to be lighter than that of wild type. These variations in fresh matter were correlated with changes in water content. The *clc-a* and *35S::NRT2.7(clc-a)* lines exhibited reduced water content compared to the wild type, while *35S::NRT2.7* showed an increase in water content (Figure 2B).

**Figure 2.**
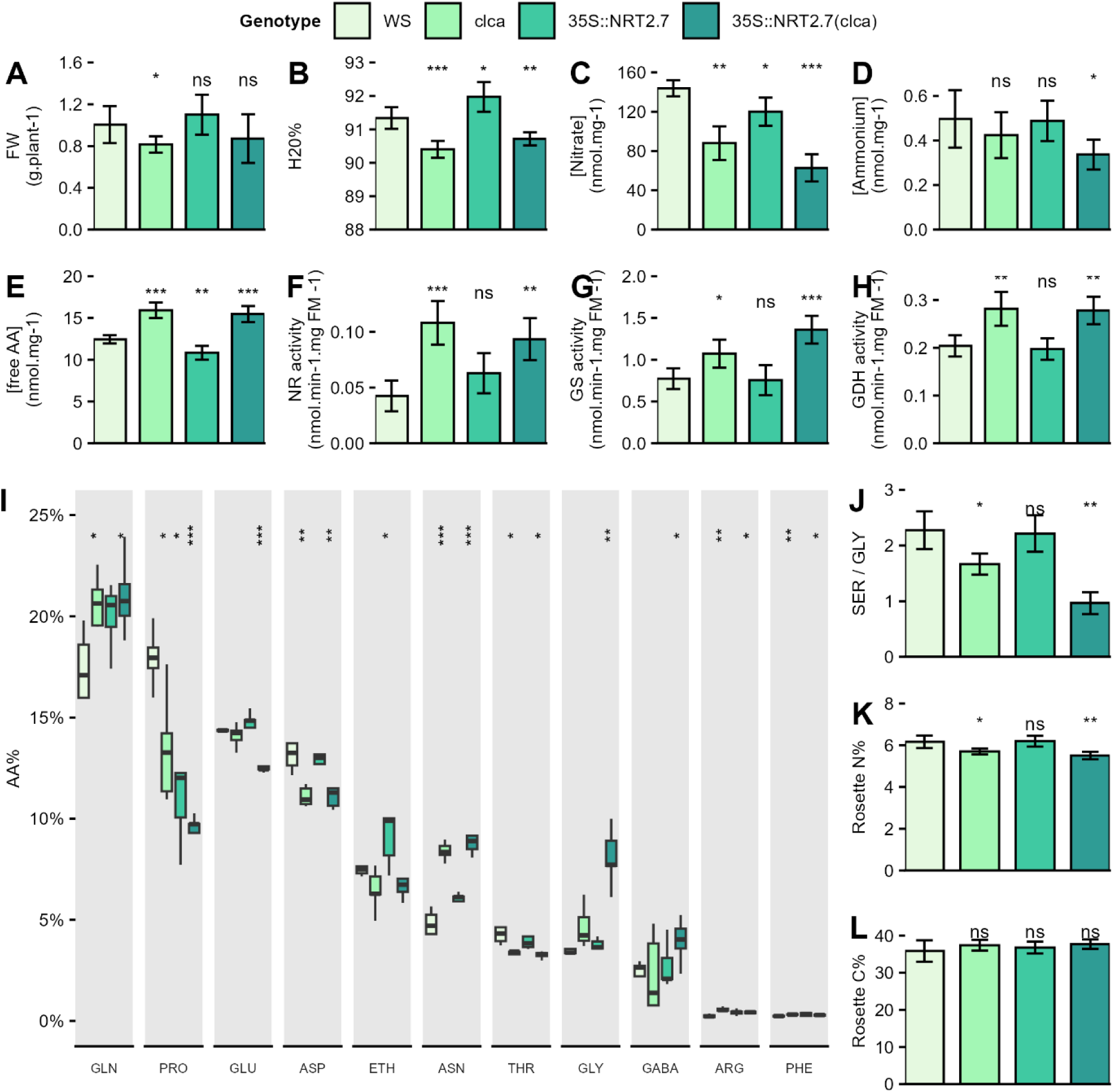
Modification of vacuolar nitrate transport decreases nitrate and ammonium pools, enhances nitrogen metabolism and increases inorganic nitrogen pools in Arabidopsis leaves. Plants were grown on sand under high-nitrate condition. Rosettes were harvested at the vegetative stage. **(A)** Rosette fresh weight (FW) per plant. **(B)** Rosette water content. **(C)** Nitrate concentration in rosettes. **(D)** Ammonium concentration in rosettes. **(E)** Concentration of total free amino acid in rosettes. **(F–H)** Maximal activities of nitrate reductase (NR), glutamine synthetase (GS), and glutamate dehydrogenase (GDH) in rosettes. **(I)** Relative abundance (as % of total amino acids) of selected free amino acids in rosettes. **(J)** SER/GLY ratio in the free amino acid pool. **(K–L)** Total nitrogen and carbon concentrations in rosettes (N% and C%). Bars represent means ± standard deviation; boxplots show the distribution of relative amino acid abundances. Stars indicate significant differences between WS and the other genotypes (t-test, *p<0.05, **p<0.01, ***p<0.001, n=6-12).

Nitrate concentration was significantly lower in *clc-a* (−43%), *35S::NRT2.7* (−14%), and *35S::NRT2.7(clc-a) (*-58%) compared to the wild type (Figure 2C). Lower nitrate concentrations was associated with lower ammonium concentrations, though only significant for *35S::NRT2.7(clc-a)*. By contrast, the concentration in free amino acids was higher in *clc-a* and *35S::NRT2.7(clc-a)* but lower in *35S::NRT2.7* compared to the wild type (Figure 2E). These results suggest that the reduced nitrate storage capacity led to greater nitrate and ammonium assimilation, driving enhanced amino acid production in *clc-a* background. Activities of key enzymes of nitrogen assimilation were modified accordingly. Nitrate reductase (NR) activity was 2.57-fold higher in *clc-a* and 2.22-fold higher in *35S::NRT2.7(clc-a)* than in the wild type (Figure 2F). Glutamine synthetase (GS) activity was 1.56-fold higher in *clc-a* and 1.75-fold higher in *35S::NRT2.7(clc-a)*, while glutamate dehydrogenase (GDH) activity was 1.38-fold and 1.36-fold higher, respectively (Figures 2J–L). In contrast, enzyme activities in *35S::NRT2.7* were similar to those in wild-type plants.

A closer look at individual amino acid profiles revealed that *clc-a* and *35S::NRT2.7(clc-a)* accumulated amino acids directly linked to nitrate and ammonium assimilation, such as glutamine, asparagine, and arginine (Figure 2I). Conversely, proline, glutamate, aspartate, ethanolamine, and threonine were less abundant in the modified lines compared to wild type. Notably, glycine levels were significantly higher in *35S::NRT2.7(clc-a)* and moderately higher in *clc-a*, generating imbalance in the serine-to-glycine ratio (Figure 2G), which suggests increased photorespiration activity in these lines.

Although amino acid pools increased, they did not compensate for the reductions in nitrate and ammonium levels. Rosette N concentration was significantly reduced by 6% in *clc-a* and 14% in *35S::NRT2.7(clc-a)* (Figure 2K). In contrast, rosette C concentration remained similar across all lines (Figure 2L), as observed in seeds (Figure 1C).

Consistent with these observations, untargeted metabolic profiling revealed genotype-dependent alterations in carbon-related metabolites (Figure S2). The *35S::NRT2.7* lines displayed a relative increase of soluble sugars, notably glucose and fructose, whereas soluble sugars were reduced in *clc-a* and 35S::NRT2.7(*clc-a*) plants. In contrast, the *35S::NRT2.7(clc-a)* line exhibited a marked enrichment in organic acids, including fumarate, succinate, citrate, and glycerate, indicating a substantial reprogramming of central carbon metabolism in response to altered vacuolar nitrate transport.

In summary, modifications in vacuolar nitrate transport enhanced nitrogen assimilation and amino acid syntheses, particularly in the *35S::NRT2.7(clc-a)* line. This line exhibited unique metabolic traits, including increased enzyme activity and altered amino acid profiles, highlighting an increase in photorespiration and nitrogen use efficiency.

### Modification of vacuolar nitrate transport affects transpiration-related traits but not stomatal density

Thermal imaging revealed significant differences in leaf temperature among the four genotypes (Figure 3A). Wild-type plants displayed the highest rosette temperatures, whereas all genotypes with modified vacuolar nitrate transport exhibited significantly cooler leaves. This was reflected by a progressive shift of the temperature distribution toward lower values in the modified lines (Figure 3B). Stomatal density did not differ significantly between genotypes (Figure 3C), indicating that changes in leaf temperature were not driven by alterations in stomatal number. In contrast, stomatal conductance (gsw) was significantly increased in the *35S::NRT2.7(clc-a)* line compared with the wild type (Figure 3D). This increase in gas exchange is consistent with enhanced transpiration and likely contributes to the observed rosette cooling in this genotype. The thermal and gas-exchange phenotypes are in line with the reduced leaf water content measured in *clc-a* and *35S::NRT2.7(clc-a)* plants (Figure 2B), supporting increased water loss in genotypes with altered vacuolar nitrate transport. Such phenotypes are consistent with the established role of CLC-A in guard cells, where disruption of vacuolar nitrate transport has been shown to affect cytosolic pH regulation, impair stomatal closure, and promote increased transpiration (Demes et al., 2020).

**Figure 3.**
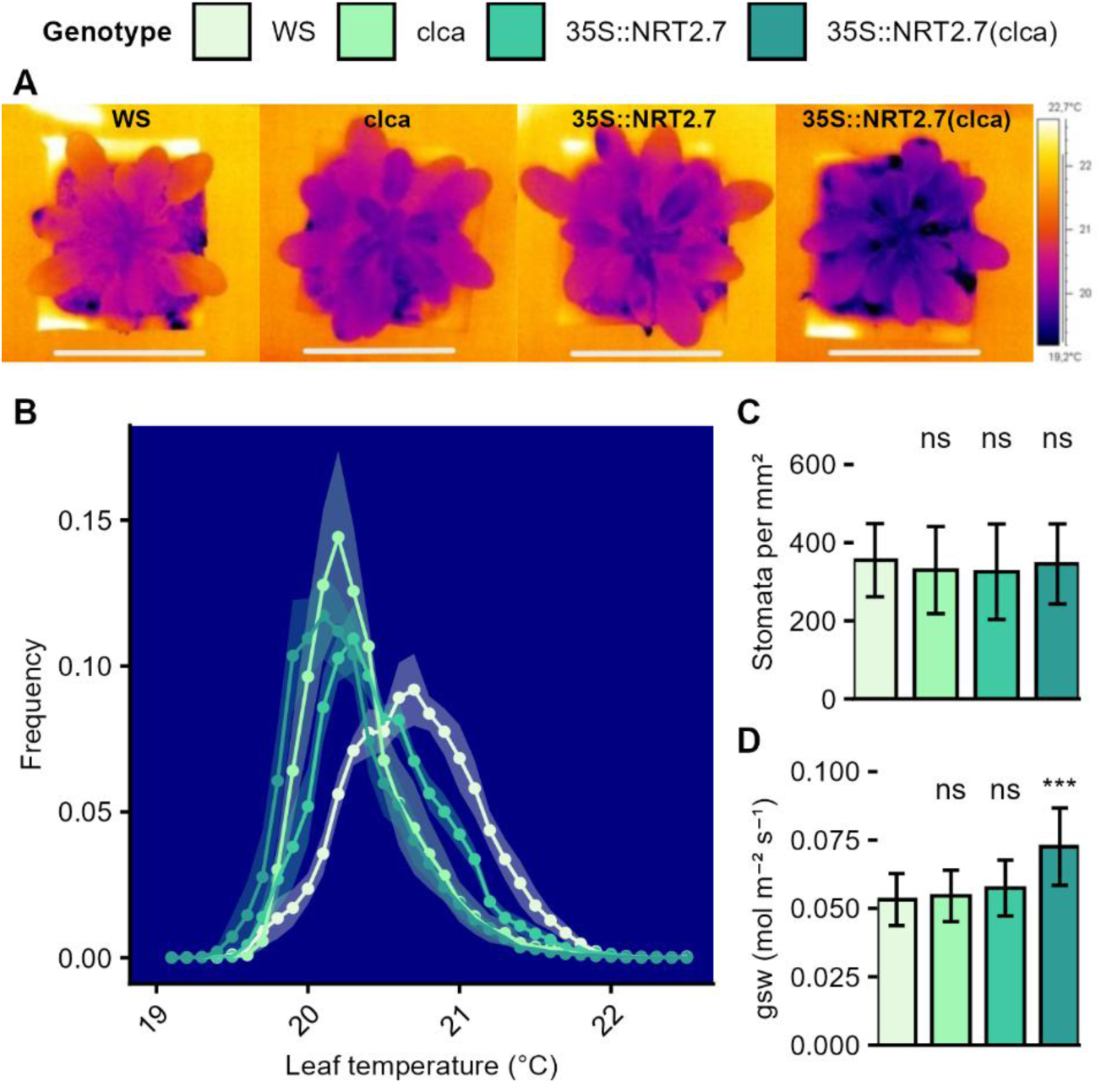
Modification of vacuolar nitrate transport affects transpiration-related traits but not stomatal density. **(A)** Representative thermal images of rosettes from the four genotypes (WS wild type, *clc-a*, *35S::NRT2.7*, and *35S::NRT2.7(clc-a)).* **(B)** Distributions of leaf temperature expressed as mean relative frequency across all plants; shaded areas indicate ± standard error. (n= 6). **(C)** Stomata density. **(D)** Stomatal conductance to water vapour (gsw) measured under controlled conditions using a LI-6800 system. Bars represent means ± standard deviation. Stars indicate significant differences between WS and the other genotypes (t-test, ***p<0.001 , n=6).

### Proteomic analysis highlights contrasting protein accumulation patterns in clc-a and NRT2.7-modified lines

Proteomic analyses were performed on vegetative leaves across the four genotypes. A total of 8,896 protein groups were identified, with 581 showing significant differences in accumulation among the genotypes. Hierarchical clustering of the differentially accumulated proteins (DAPs) revealed five distinct groups, each enriched in specific GO terms (Figure 4). Notably, the *clc-a* mutant and the *35S::NRT2.7* overexpressing line exhibited opposite profiles in protein abundance patterns. Interestingly the *35S::NRT2.7(clc-a)* displayed a combined pattern that integrated features of both single modifications along with specific alterations enlighten by the Group 4 proteins.

**Figure 4.**
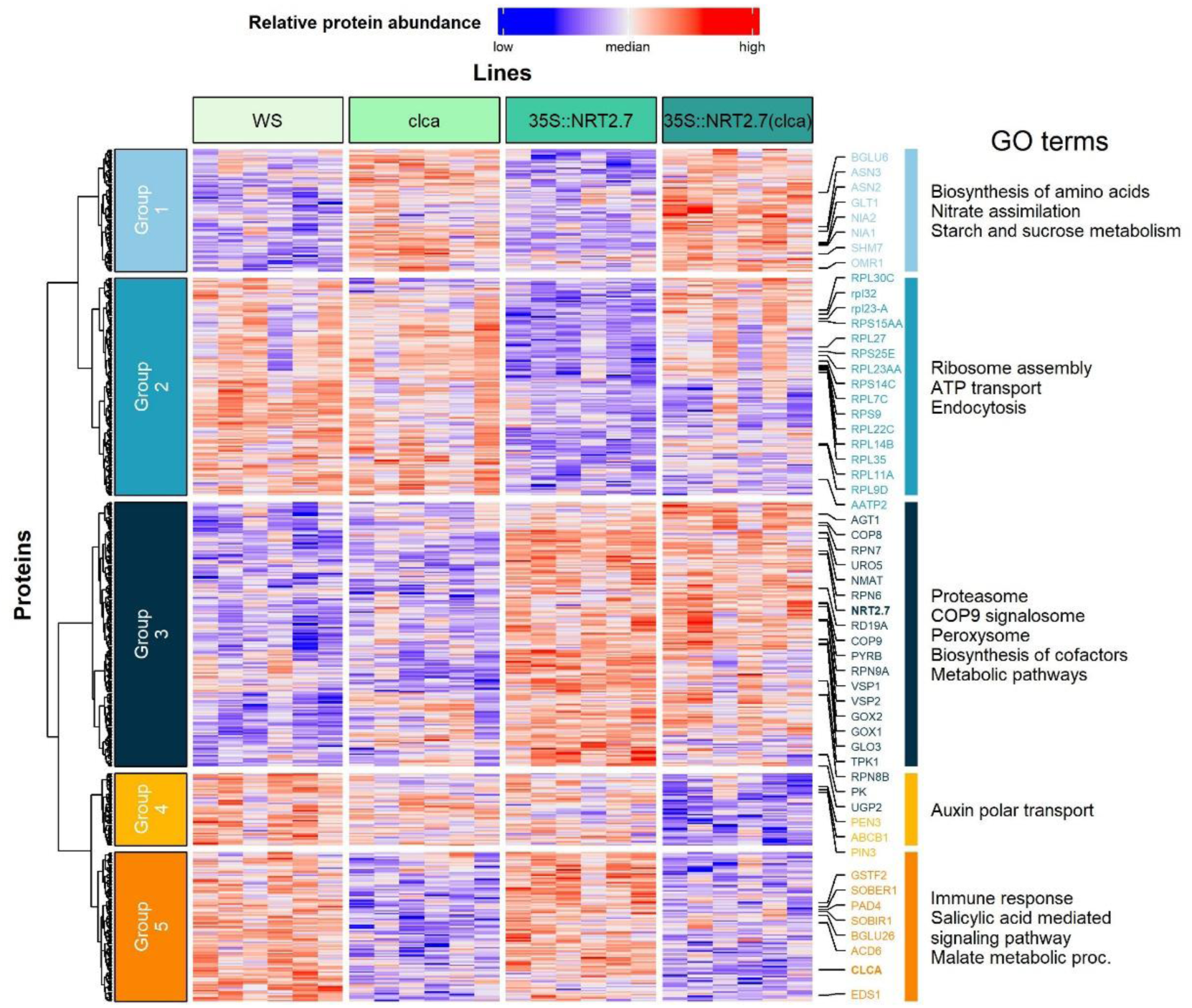
Modification of vacuolar nitrate transport modifies proteome in Arabidopsis leaves. Heatmap of differentially abundant proteins identified in rosettes of WS, *clc-a*, *35S::NRT2.7* and *35S::NRT2.7(clc-a)* plants (six biological replicates per genotype). Proteins showing a significant genotype effect (n=6, one-way ANOVA, p<0.01) were clustered using Ward’s method and grouped into five major clusters (left blocks). Representative GO terms associated with each cluster are shown on the right. Genotypes are indicated by color blocks at the top. Selected proteins related to nitrate transport and assimilation, amino acid metabolism, translation, proteasome/COP9 signaling, auxin transport and defense responses are annotated. Color scale represents relative protein abundance (low in blue to high in red).

Group 1 comprised proteins that were more abundant in both *clc-a* and *35S::NRT2.7(clc-a)* lines compared to the wild type and *35S::NRT2.7*. This group was enriched in proteins involved in amino acid biosynthesis, nitrate assimilation, and carbohydrate metabolism, including NIA1, NIA2, ASN2, and GLT1. The enrichment of these proteins is consistent with an enhanced nitrogen assimilation capacity in the *clc-a* background. Group 2 included proteins that were predominantly less abundant in the *35S::NRT2.7* line and, to a lesser extent, in the *35S::NRT2.7(clc-a)* line. These proteins were mainly associated with ribosomal function, ATP transport, and endocytosis. The coordinated decrease of these functional categories suggests a reduction in translational and cellular transport processes in *NRT2.7* overexpressing lines. Group 3 consisted of proteins that accumulated preferentially in the NRT2.7 overexpression backgrounds, namely 35S::NRT2.7 and 35S::NRT2.7(*clc-a*). This group included NRT2.7 itself, nitrogen storage proteins (VSP1, VSP2), components of the proteasome (RPN6, RPN7, RPN8.B, RPN9.A), COP9 signalosome subunits (COP8, COP9), and several peroxisome-associated proteins involved in photorespiratory and metabolic processes (PK, GOX1, GOX2, GLO3). The enrichment of these proteins is indicative of enhanced metabolic turnover and protein degradation pathways in *NRT2.7* overexpressing lines. Group 4 represented a set of protein abundance changes that were specific to the double mutant 35S::NRT2.7(*clc-a*). This group was enriched in proteins involved in auxin polar transport, including PEN3, ABCB1, and PIN3, which were not similarly affected in either single genotype. These coordinated changes in auxin transport-related proteins suggest a specific alteration of auxin distribution or signaling in the double mutant. Group 5 contained proteins whose abundance was reduced in the *clc-a* mutant and in the 35S::NRT2.7(*clc-a*) line compared to the wild type and 35S::NRT2.7. These proteins were mainly associated with salicylic acid– related pathways and immune responses, including PAD4, ACD6, SOBIR1, and BGLU26. Their reduced abundance is consistent with an attenuation of salicylic acid– associated defense pathways in the *clc-a* background.

Overall, the proteomic analysis revealed genotype-specific and combined effects of vacuolar nitrate transport modifications on protein accumulation patterns. The *clc-a* background was characterized by increased abundance of proteins involved in nitrogen and carbon metabolism together with reduced abundance of immune-related proteins, whereas *35S::NRT2.7* lines showed enrichment in storage proteins, proteasome components, and peroxisomal metabolism. The double mutant *35S::NRT2.7(clc-a)* displayed an intermediate and integrative profile, combining features of both parental genotypes while exhibiting unique alterations, notably in proteins related to auxin transport and signaling.

### Modifications of vacuolar nitrate transport improve nitrogen remobilization and nitrogen use efficiency for seed filling

We conducted a ^15^N nitrate labeling experiment across two independent cultures to determine how combining *clc-a* and *35S::NRT2.7* affects nitrate uptake and nitrogen remobilization.

Despite reduced rosette nitrogen concentration in the modified lines compared to the wild type (Figure 5A; Figure 2K), no significant differences were observed between genotypes for nitrate uptake during the vegetative stage. Uptake rates were approximately 0.12 mg N day^-1^ across all lines (Figure 5B). This lack of difference likely reflects the early developmental stage of the plants, characterized by limited rosette expansion and relatively low nitrogen demand. Nitrate uptake was subsequently monitored at five time points during the reproductive phase (7, 14, 21, 28, and 35 days after emergence of the flowering bud, dae). Nitrate uptake decreased progressively over time in all genotypes, following a similar overall trajectory (Figure 5C). However, at 14 dae, nitrate uptake was significantly lower in all modified lines (*clc-a*, *35S::NRT2.7*, and *35S::NRT2.7(clc-a)*) compared with the wild type. At this stage, nitrate uptake was 0.75 mg N day^-1^ in wild type, while *clc-a*, *35S::NRT2.7*, and *35S::NRT2.7(clc-a)* nitrate uptake rates were 0.61, 0.55, and 0.58 mg N day^-1^, respectively. By 35 dae, the nitrate uptake rates converged to approximately 0.39 mg N day^-1^ in all the genotypes (Figure 5C). These results indicate that the decline in nitrate uptake occurred earlier during the reproductive phase in the modified lines than in the wild type.

**Figure 5.**
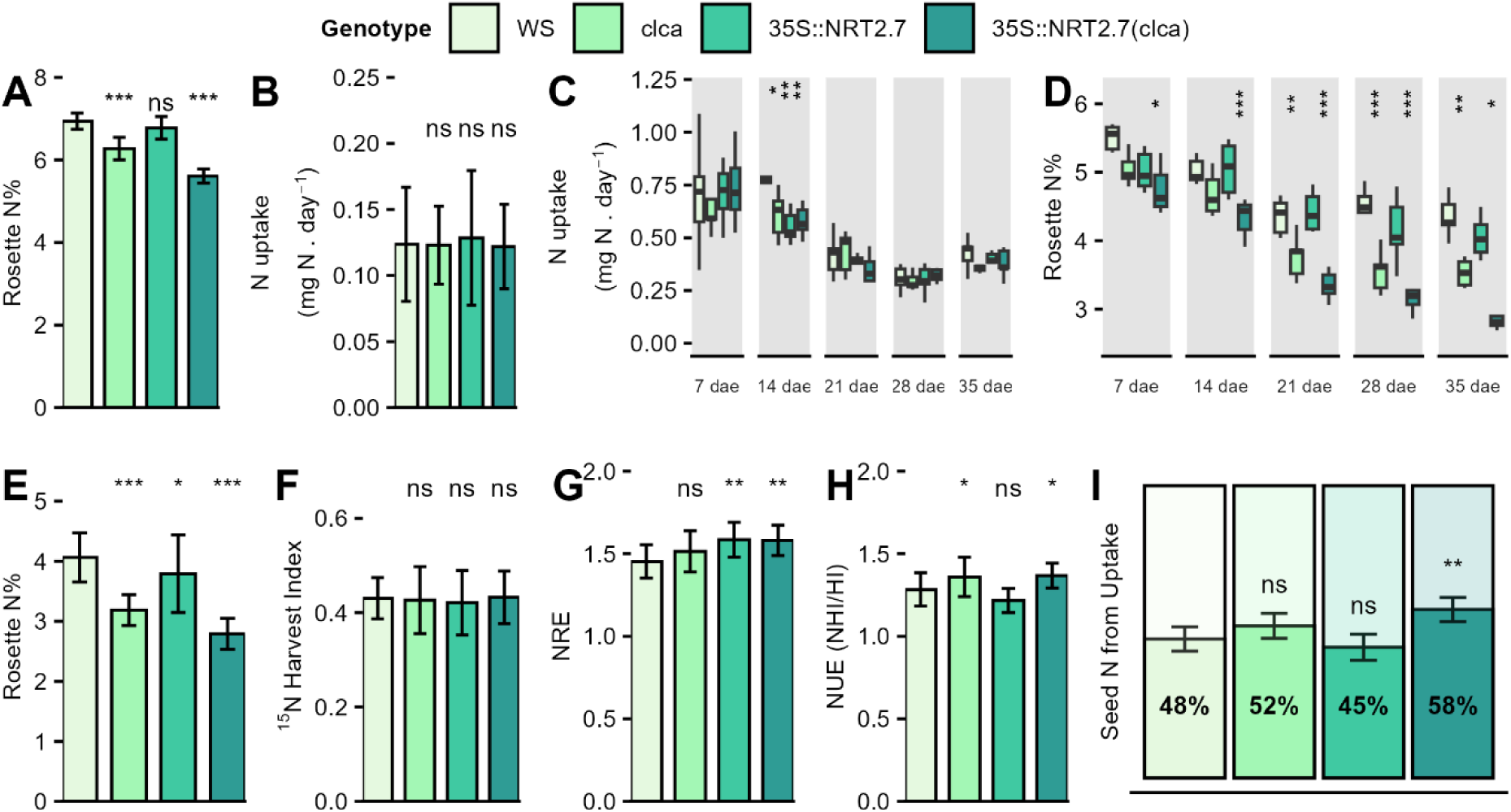
Modification of vacuolar nitrate transport increases nitrogen mobilization efficiency and N uptake to the seeds. Plants were grown on sand under high-nitrate conditions. Vegetative nitrate uptake was quantified in rosettes during a short-term ^15^N-labelling experiment, and reproductive uptake and remobilization were analyzed in plants labelled repeatedly with ^15^NO_3_ during the reproductive phase. **(A)** Rosette nitrogen concentration at the vegetative stage. **(B)** Vegetative nitrogen uptake. **(C)** Reproductive nitrogen uptake at successive time points (dates after emergence). **(D)** Rosette nitrogen concentration. **(E)** Rosette nitrogen concentration (N%) at harvest in the remobilization experiment. **(F)** ^15^N harvest index (^15^NHI) **(G)** Nitrogen remobilization efficiency calculated as ^15^NHI/HI **(H)** Nitrogen use efficiency calculated as NHI/HI **(I)** Fraction of seed-nitrogen originating from post-labelling uptake. Bars represent means ± standard deviation; boxplots (C, D) show the distribution of individual plants. Stars indicate significant differences between WS and the other genotypes (t-tests, *p<0.05, **p<0.01, ***p<0.001, n=16).

Nitrogen concentration measurements performed on the same plants across the five time points provided additional insight. By 35 dae, nitrogen concentration in rosettes had declined more sharply in the modified lines than in the wild type (Figure 5D). The reduction amounted to 20% in wild type, 30% in *clc-a*, 20% in *35S::NRT2.7*, and 33% in *35S::NRT2.7(clc-a).* The measurement of nitrogen concentrations in the remains of the dry rosette harvested at seed maturity in the remobilization experiment (Figure 5E) showed interestingly the same pattern as the 35 dae rosettes (Figure 5D). At this stage, rosette N concentrations were significantly lower in all the modified lines, particularly in those carrying the *clc-a* mutation. Because rosette nitrogen concentration reflects the dynamic balance between nitrogen influx (uptake) and efflux (remobilization), the stronger decline observed in the modified lines suggests either reduced post-flowering nitrogen uptake or enhanced nitrogen remobilization from rosette leaves, or both, compared with the wild type.

To directly assess nitrogen remobilization from vegetative organs to the seeds, we calculated the ^15^N Harvest Index (^15^NHI; i.e. ^15^N partition in seeds) following pulse– chase labeling experiments, as described by Marmagne et al. (2020). Briefly, plants were labeled with ^15^NO₃⁻ at the vegetative stage, and ^15^N content was subsequently quantified in individual organs at maturity, allowing us to calculate the partition of ^15^N in different organs at the end of the plant cycle, and thus track the translocation of the ^15^N that have been taken up by the plant at vegetative stage. The proportion of ^15^N allocated to seeds (^15^NHI; Figure 5F) did not differ significantly between genotypes, indicating that independently of the biomass of the different source and sink organs, the same proportion of ^15^N was translocated from vegetative tissues to the seeds irrespective of genotype. However, when normalized to the harvest index (HI), which reflects the relative dry biomass of sink and source compartments, the ratio of ^15^NHI to HI (hereafter referred to as nitrogen remobilization efficiency; NRE) was higher in the modified lines than in the wild type. Higher NRE in *35S::NRT2.7* and *35S::NRT2.7(clc-a)* was significant and consistent tendency was observed in *clc-a* (Figure 5G).

In conclusion, modifications of vacuolar nitrate transport did not affect nitrate uptake during the vegetative phase but led to an earlier decline in nitrate uptake during reproductive development and to enhanced nitrogen remobilization efficiency. These effects, most pronounced in the 35S::NRT2.7(clc-a) line, contribute to improved nitrogen use efficiency (Figures 5H). Importantly, analysis of the origin of seed nitrogen further showed that the relative contribution of post-flowering nitrogen uptake to seed nitrogen content differed among genotypes (Figure 5I). This contribution accounted for approximately 48% in the wild type, 52% in *clc-a*, 45% in *35S::NRT2.7*, and reached 58% in the *35S::NRT2.7(clc-a)* line. Together, these results indicate that the double mutant combines enhanced nitrogen remobilization efficiency with an increased contribution of post-flowering nitrogen uptake to seed filling, thereby maximizing nitrogen allocation to seeds and contributing to improved overall nitrogen use efficiency.

### *NRT2.7* and its *HvNRT2.10* barley homologue play the same role in nitrogen use efficiency

Nitrogen remobilization during grain filling is a key agronomic trait in cereals, as it strongly influences grain nitrogen content and overall nitrogen use efficiency. To assess whether the role of NRT2.7 is conserved across plant species, we cloned the barley homolog *HvNRT2.10* and expressed it under the control of the 35S promoter in Arabidopsis wild-type and *clc-a* mutant backgrounds.

Across two independent cultures, *HvNRT2.10*-overexpressing lines displayed reduced nitrogen concentrations in rosette tissues compared with the wild type. On average, rosette nitrogen concentration decreased by 24% in the two *35S::HvNRT2.10* lines and by 35% in the two *35S::HvNRT2.10(clc-a)* lines (Figure 5A). In parallel, seed nitrogen concentration increased in the overexpressing lines, with mean increases of 6% in *35S::HvNRT2.10* and 12% in *35S::HvNRT2.10(clc-a)* relative to the wild type (Figure 5B).

The partitioning of labeled nitrogen to seeds, expressed as the ^15^N Harvest Index (^15^NHI), did not differ significantly among genotypes (average 0.40; Figure 5C), showing that *HvNRT2.10* overexpression did not play prominent role in nitrogen remobilization. indicating that *HvNRT2.10* overexpression did not markedly alter the overall capacity for nitrogen remobilization from vegetative tissues to seeds. However, when normalized to the harvest index, NRE was significantly higher in the *HvNRT2.10*-overexpressing lines. NRE increased by 12% in *35S::HvNRT2.10* and by 23% in the *35S::HvNRT2.10(clc-a)* lines (Figure 5D), revealing a more efficient use of remobilized nitrogen relative to sink size, particularly in the double mutant.

Consistent with these changes, NUE was also improved, with average increases of 11% in *35S::HvNRT2.10* and 22% in *35S::HvNRT2.10(clc-a)*. These responses closely mirror those observed in the corresponding Arabidopsis lines overexpressing AtNRT2.7, either alone or in the clc-a background (Figures 4F–H). Together, these results demonstrate that HvNRT2.10 can functionally substitute for AtNRT2.7 in Arabidopsis, supporting a conserved role of NRT2.7-related transporters in enhancing nitrogen remobilization efficiency and nitrogen use efficiency, with the strongest effects observed when combined with reduced vacuolar nitrate storage.

## DISCUSSION

Recent studies have highlighted the central role of vacuolar nitrate fluxes in regulating whole-plant nitrogen homeostasis and improving NUE, either by reducing nitrate storage in the vacuole or by increasing its mobilization toward metabolic and growth processes (Han et al., 2016; Lu et al., 2022; Hodin et al., 2023; Armengaud et al., 2025). These findings have renewed interest in the functional characterization of vacuolar nitrate transporters and in how their activities shape nitrogen allocation between vegetative and reproductive organs. Here, we combined the disruption of CLC-A, the major vacuolar nitrate importer in leaves, with the overexpression of NRT2.7, a tonoplast transporter previously associated with seed nitrate accumulation and recently proposed to facilitate nitrate efflux. By integrating physiological, metabolic, and proteomic analyses, we show that genetically modifying vacuolar nitrate storage and export profoundly alters nitrogen metabolism and enhances nitrogen remobilization toward seeds and overall NUE. Notably, the 35S::NRT2.7(clc-a) line displayed cumulative and unique metabolic features, indicating that CLC-A and NRT2.7 act through distinct yet complementary mechanisms to control vacuolar nitrate dynamics and whole-plant nitrogen allocation.

### Seed protein content increases without carbon penalties

A central and unexpected finding of this study is that modifying vacuolar nitrate transport substantially increases seed protein concentration without detectable penalties on seed carbon content, lipid amount, or lipid composition (Figure 1). This result is striking because it challenges the well-established negative relationship between nitrogen-and carbon-based storage compounds in seeds. In Arabidopsis (Marmagne et al., 2020), as well as in major oilseed crops such as sunflower (Li et al., 2024), rapeseed (Gunasekera et al., 2006; Peltonen-Sainio et al., 2011), and soybean (Morrison et al., 2000), increases in seed protein content typically occur at the expense of lipid accumulation. This negative association is remarkably robust across genotypes and environmental conditions. Marmagne et al. (2020) showed that the inverse relationship between seed N% and C% is maintained even when nitrogen remobilization efficiency varies strongly or when post-flowering stresses alter source-sink dynamics. From a metabolic perspective, increased protein synthesis imposes substantial energetic and reductant costs associated with nitrate uptake, reduction, and amino acid biosynthesis, which are generally compensated by reduced carbon allocation to storage compounds (Munier-Jolain and Salon, 2005). Similar negative relationships between grain protein concentration and yield are well documented in cereals (Oury and Godin, 2007; Laidig et al., 2017; Geyer et al., 2022). Our results therefore indicate that seed protein content can be increased without compromising carbon deposition or yield, a long-standing objective in crop improvement. By alleviating a fundamental constraint on seed C:N stoichiometry, manipulation of vacuolar nitrate transport emerges as a promising strategy to improve seed nutritional quality while preserving carbon-based yield components.

### Complementary roles of CLC-a and NRT2.7 in whole-plant nitrogen allocation

Our results indicate that altering vacuolar nitrate transport at different steps of the pathway differentially affects nitrogen allocation between vegetative and reproductive organs. Disruption of *CLC-A* primarily impacted nitrogen partitioning in source tissues, whereas overexpression of *NRT2.7* mainly affected nitrogen retention in seeds (Figure 5), revealing complementary roles for these transporters. Consistent with previous studies identifying CLC-A as a major regulator of vacuolar nitrate storage in leaves (De Angeli et al., 2006; Monachello et al., 2009; Hodin et al., 2023), the *clc-a* mutant displayed reduced nitrogen retention in rosettes during reproductive development and enhanced nitrogen transfer toward reproductive organs. This supports the view that limiting vacuolar nitrate buffering in vegetative tissues facilitates nitrogen remobilization from source leaves and improves NUE despite lower nitrate pools. In contrast, overexpression of *NRT2.7* preferentially modified nitrogen allocation at the sink level. In agreement with its role as a tonoplast-localized transporter strongly expressed in seeds (Chopin et al., 2007; David et al., 2014), *NRT2.7* overexpression increased seed nitrate and nitrogen concentrations without substantially altering vegetative nitrogen pools (Figures 1, 2). This suggests that NRT2.7 primarily reinforces seed sink strength rather than promoting nitrogen export from source tissues. Importantly, the combined modification in the *35S::NRT2.7(clc-a)* line integrated both effects, resulting in reduced nitrogen retention in vegetative tissues together with enhanced nitrogen accumulation in seeds (Figure 5). This dual phenotype led to the highest seed nitrogen and protein content among the genotypes analyzed (Figure 1), indicating that vacuolar nitrate storage in leaves and nitrate loading in seeds represent partially independent but complementary regulatory layers controlling whole-plant nitrogen allocation.

### Metabolic reprogramming in the rosette under altered vacuolar nitrate flux

Modifications of vacuolar nitrate transport were associated with a profound reprogramming of nitrogen and carbon metabolism in vegetative tissues. In genotypes with reduced vacuolar nitrate buffering, particularly the *clc-a* mutant and the *35S::NRT2.7(clc-a)* line, enhanced activities of key nitrate assimilation enzymes indicate an increased capacity to convert inorganic nitrogen into organic forms (Figures 2F,G,H). This enzymatic signature is consistent with reduced nitrate and ammonium pools in rosettes and supports the view that limiting vacuolar nitrate storage shifts nitrogen metabolism toward rapid assimilation rather than inorganic nitrogen accumulation. This acceleration of nitrogen assimilation was accompanied by changes in amino acid composition (Figures 2I, 4), including increased pools of glutamine, asparagine, and arginine, which are central intermediates for nitrogen transport and temporary storage (Masclaux-Daubresse et al., 2010). Importantly, these metabolic changes occurred without an increase in total rosette nitrogen content (Figures 2K, 5A, 5E), indicating altered nitrogen turnover and remobilization dynamics rather than enhanced nitrogen uptake. In parallel, several indicators point to a coordinated adjustment of carbon metabolism. The elevated Gly/Ser ratio observed in clc-a and in the double mutant (Figure 2J) suggests enhanced photorespiratory activity, a process tightly interconnected with both carbon and nitrogen metabolism (Shi and Bloom, 2021). Consistently, proteomic analyses revealed increased abundance of enzymes involved in amino acid biosynthesis, glycolysis, the tricarboxylic acid cycle, and peroxisomal reactions, particularly in the clc-a background and in the 35S::NRT2.7(clc-a) line (Figure 4).

The schematic synthesis presented in Figure 7 integrates these metabolic and proteomic observations into a unified framework. Reduced nitrate sequestration in the vacuole increases cytosolic nitrate availability, thereby stimulating nitrate assimilation in the chloroplast. This enhanced nitrogen flux is supported by the activation of glycolysis and the tricarboxylic acid cycle - which provide carbon skeletons and reducing power - and by the increase of photorespiratory activity, which may act as a metabolic hub coordinating carbon and nitrogen fluxes. Importantly, these adjustments maintain carbon metabolism, as evidenced by stable sugar levels and unchanged carbon content, despite increased nitrogen assimilation and amino acid biosynthesis. Classical energetic models of plant nitrogen metabolism indicate that nitrate assimilation represents a major metabolic cost, requiring substantial investments of carbon skeletons, ATP, and reducing equivalents (Gutschick, 1981). In this context, the coordinated upregulation of nitrate assimilation enzymes, central carbon metabolism, and photorespiration observed in the present study is consistent with an increased cellular energy demand, particularly in the 35S::NRT2.7(*clc-a*) line. Although direct measurements of energy charge or respiratory activity were not performed, the convergence of metabolic, enzymatic, and proteomic signatures supports the hypothesis that altered vacuolar nitrate fluxes place a strong demand on cellular energy metabolism, enabling enhanced nitrogen assimilation and redistribution while preserving carbon homeostasis.

### Conservation of the mechanism in cereals and implications for crop improvement

The translational relevance of modifying vacuolar nitrate transport is supported by the functional conservation of NRT2.7-related mechanisms. Heterologous expression of the barley homolog *HvNRT2.10* in Arabidopsis reproduced key features of the *AtNRT2.7* overexpression phenotype, including enhanced nitrogen allocation to seeds and improved nitrogen remobilization efficiency, particularly in the *clc-a* background (Figure 6). This provides proof of concept for cross-species conservation of this regulatory mechanism. Consistent with this view, *HvNRT2.10* shows high expression in leaves and substantial expression in seeds in barley (Guo et al., 2020; Zoghbi-Rodríguez et al., 2021), is induced under nitrogen limitation, and has been identified as a candidate gene associated with NUE in genome-wide association studies(Guo et al., 2020; Karunarathne et al., 2020; Decouard et al., 2022). Together, these observations are consistent with a role for HvNRT2.10 in nitrogen redistribution rather than in primary root nitrate uptake. Similar conclusions emerge from studies in wheat, where members of the *NRT2* family have been associated with grain nitrogen accumulation independently of yield, notably through the trait of grain protein deviation (GPD). In durum wheat, *TdNRT2-6A*, a homolog of *AtNRT2.7*, was significantly associated with GPD in a genome-wide association study, identifying this locus as a determinant of grain protein content uncoupled from yield penalties (Nigro et al., 2019). This conclusion was further strengthened by the identification of *TaNRT2.6-7A*, the bread wheat homolog of *AtNRT2.7*, as a strong candidate gene underlying a major GPD quantitative trait locus (Geyer et al., 2022). Notably, *TaNRT2.6-7A* is predominantly expressed in leaves throughout development, whereas most other wheat NRT2 genes are primarily root-expressed, supporting a conserved role of the leaf-expressed *NRT2* transporters in nitrogen redistribution toward developing grains.

**Figure 6.**
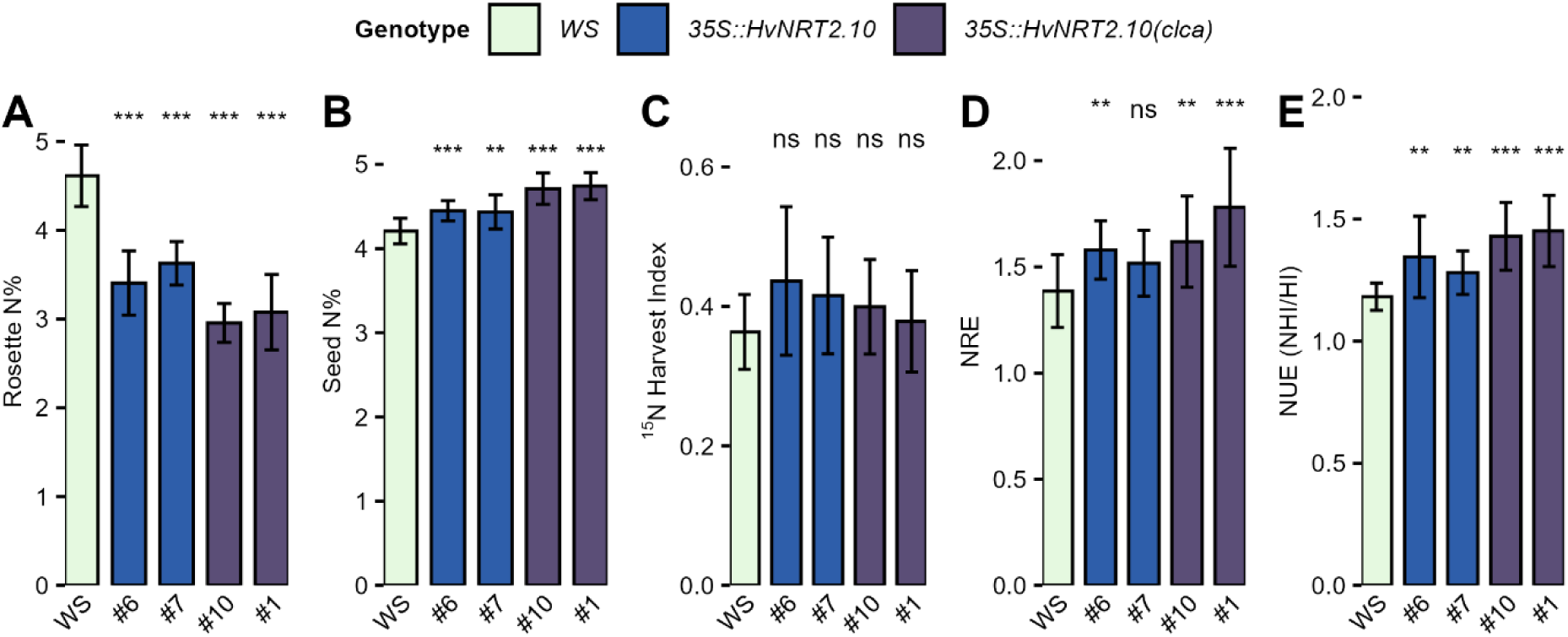
Modification of vacuolar nitrate transport using *35S::HvNRT2.10* shows the conserved role of NRT2.7 barley homologue on NUE in Arabidopsis and barley. Plants were grown under high-nitrate condition and labelled with 10 mM K^15^NO₃ to quantify nitrogen partitioning and fluxes. Nitrogen traits were measured on Arabidopsis wild type and *clc-a* mutant expressing *HvNRT2.10* under the control of the 35S constitutive promoter. **(A)** rosette nitrogen concentration, **(B)** seed nitrogen concentration, **(C)** ^15^N Harvest Index (^15^NHI), **(D)** Nitrogen Remobilization Efficiency (NRE = ^15^NHI/HI), **(E)** Nitrogen Use Efficiency (NUE = NHI/HI) . Bars show means ± standard deviation. Stars indicate significant differences relative to wild type (t-test , *p<0.05, **p<0.01, ***p<0.001, n=12).

**Figure 7.**
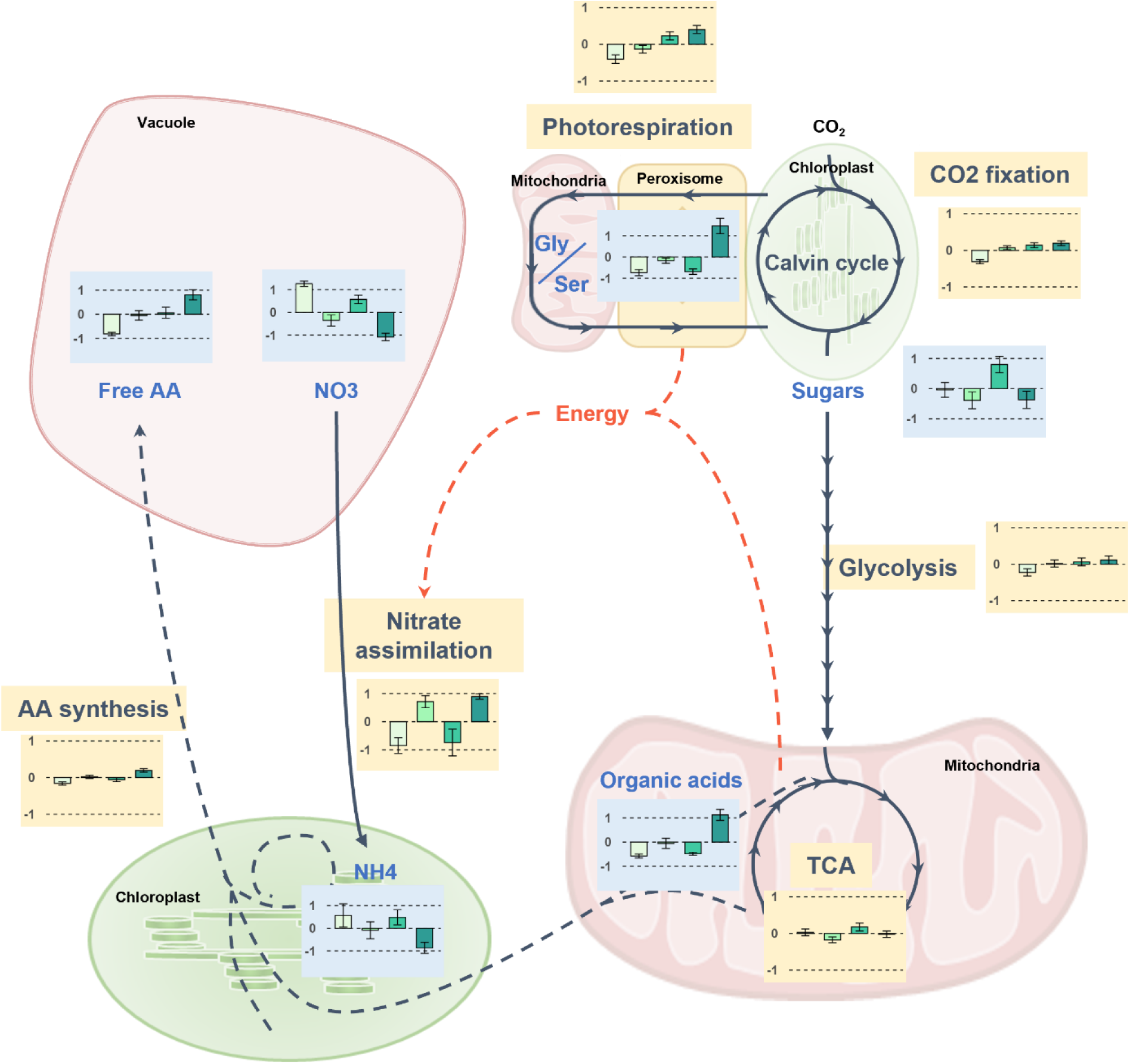
Schematic summary of the effects of modification of vacuolar nitrate transport on nitrogen metabolism. Blue bar plots represent the relative metabolite contents (free amino acids, nitrate, ammonium, sugars, organic acids and Gly/Ser ratio). Green bar plots represent the relative abundance of proteins involved in TCA cycle, nitrate assimilation, amino acid biosynthesis, glycosis, CO_2_ fixation and photorespiration.

Beyond nitrate uptake systems, vacuolar nitrate storage mechanisms also appear to be conserved in cereals. Homologs of CLC-A are present in wheat and other cereal genomes and have been implicated in vacuolar anion homeostasis, nitrate accumulation, and stress responses (Kumar et al., 2022). However, although these studies clearly establish a role for cereal CLC proteins in vacuolar nitrate storage and ion balance, their specific contribution to nitrogen remobilization efficiency and grain nitrogen allocation has not yet been directly demonstrated in these species. Within this context, the Arabidopsis data presented here provide a mechanistic framework linking vacuolar nitrate compartmentation to nitrogen remobilization efficiency and seed nitrogen allocation. The enhanced phenotypes observed when reduced vacuolar nitrate storage is combined with increased nitrate flux toward seeds, as exemplified by the *clc-a* background, show that vacuolar nitrate buffering acts as a regulatory node controlling the timing and efficiency of nitrogen assimilation and redistribution. The fact that cereal *NRT2* family members associated with GPD are preferentially expressed in leaves, together with the conservation of *CLC-A* homologs involved in nitrate storage, argues for the existence of a comparable regulatory architecture in cereals, even though functional validation remains to be established.

From an applied perspective, our work highlights vacuolar nitrate transporters as promising targets for crop improvement that deserve attention. Unlike strategies aimed at increasing nitrogen uptake per se, modulating nitrate storage and fluxes within source tissues offers a means to enhance nitrogen use efficiency and grain protein content without necessarily exacerbating the long-standing trade-off between yield and quality. Future work combining genetic, physiological, and field-based approaches in cereals will be required to test whether coordinated manipulation of vacuolar nitrate storage and leaf-to-grain nitrogen flux can reproduce the beneficial outcomes observed in Arabidopsis and thereby contribute to more sustainable nitrogen management in crop production systems.

## EXPERIMENTAL PROCEDURES

### Plant material

All *Arabidopsis thaliana* genotypes used in this study were obtained or generated in the *Wassilewskija* (Ws) background (Armengaud et al., 2025). The *clc-a-2* T-DNA knockout mutant was sourced from the Versailles Arabidopsis Stock Center (FST 171A06, line EAE89) (Ricou et al., 2024) and is referred to as *clc-a* throughout the manuscript. The *NRT2.7* overexpression line previously described by Chopin et al. (2007) is referred to as *35S::NRT2.7*. The line overexpressing *NRT2*.7 in the *clc-a* background, generated by crossing *35S::NRT2.7* with the *clc-a-2* mutant, is referred to as *35S::NRT2.7(clc-a)*.

For heterologous expression studies, the full-length coding sequence of *HvNRT2.10* (*HORVU.MOREX.r3.7HG0729020*), from the start codon to the stop codon, was synthesized by GeneArt Life Technologies (Thermo Fisher Scientific, Germany) and cloned into the pMK-RQ vector. The CDS was subsequently transferred into the pDONR207 entry vector using a BP clonase reaction, and then recombined into the binary destination vector pMDC32 using Gateway™ technology (Thermo Fisher Scientific), yielding the *35S::HvNRT2.10* construct. Plant transformation was performed by *Agrobacterium tumefaciens* strain GV3101 (pMP90) using the floral dip method (Clough and Bent, 1998). The *35S::HvNRT2.10* construct was introduced into both *Ws* and *clc-a* backgrounds. For each genetic background, two independent homozygous single-insertion T3 lines were selected for further analyses.

### Growth conditions

For seed production, plants were grown on sand according to Masclaux-Daubresse and Chardon (2011) in a controlled-environment growth chamber (21/17 °C day/night; 160 µmol m⁻² s⁻¹) under high-nitrate conditions (10 mM nitrate). Plants were cultivated in short days (8 h light/16 h dark) for 50 days after sowing (das) and then transferred to long-day conditions (16 h light/8 h dark) to induce flowering. For rosette production used in metabolic and proteomic analyses, plants were grown on sand under high-nitrate conditions in short days for 50 days.

### Seeds analyses

#### Seed weight

Pools of 100 seeds were dried at 70 °C for 24 h and weighed using a microbalance (XS3DU, Mettler Toledo, Viroflay, France).

#### Protein dosage

Protein dosage from 100 dry seeds was performed as described by Di Berardino et al. (2018).

#### SDS-PAGE electrophoresis

Protein extraction from 100 dry seeds, SDS–PAGE separation, Coomassie staining, and western blotting were performed as described in Deruyffelaere et al. (2015). Protein band intensity was quantified using ImageJ (https://imagej.nih.gov/ij/).

#### Analysis of nitrate

Seed nitrate content was measured on aqueous extracts prepared from 2 mg of dry seeds using a spectrophotometric method adapted from Miranda et al. (2001).

#### Fatty acid Content and composition

Total fatty acids were extracted and quantified from dry seeds according to Li et al. (2006).

### Rosette analyses

#### Metabolite Extraction and Analysis

Amino acids and ammonium (Rosen, 1957) from rosette were determined as described by Diaz et al. (2005). For nitrate determination, 5 mg of fresh rosette powder were extracted using a three-step ethanol-water procedure as described by (Loudet et al., 2003). Extracts were evaporated, resuspended in water, and analysed for nitrate content according to Miranda et al. (Miranda et al., 2001).

#### Enzymatic Assays

Enzymes were extracted from frozen leaf material stored at −80 °C. Maximal extractable nitrate reductase activity was assayed following Ferrario-Méry (1997). Glutamine synthetase (GS) activity was determined according to O’Neal and Joy (1973). NADH^-^ and NAD^+^-dependent glutamate dehydrogenase activities were measured as described by Turano et al.(1996), except that the extraction buffer was identical to that used for GS.

#### Water percentage

The water percentage of rosettes was calculated from the fresh and dry weights of ground rosette material. A known quantity of freshly ground tissue was dried at 70 °C for 48 h and weighed using a microbalance (XS3DU, Mettler Toledo, Viroflay, France).

#### Infrared thermography

Thermal images were acquired with a FLIR A320 infrared camera positioned 30 cm above the rosette and controlled using ThermaCAM Researcher Pro 2.9 software. Radiometric data were exported as CSV files containing the temperature of each pixel within the rosette. Pixel temperatures were processed in R: for each plant, all pixel values were imported, cleaned, and used to calculate the relative frequency of each temperature class. Relative frequencies were then averaged across six plants of each genotype to obtain genotype-level temperature distributions and their standard errors. These distributions were plotted as mean frequency curves. Representative thermal images for each genotype were exported as JPEG files and assembled using the *cowplot* and *gridExtra* packages.

#### Stomatal conductance

Stomatal conductance to water vapour (gsw) was measured using a LI-6800 infrared gas analyser (LI-COR Biosciences). Measurements were performed between 09:00 and 12:00 on fully expanded rosette leaves under controlled conditions (150 µmol photons m⁻² s⁻¹ irradiance, 400 ppm CO₂, leaf temperature ∼27 °C, vapour pressure deficit ∼1.5 kPa).

#### Stomatal Density

Stomatal density was measured on leaves collected at the same developmental stage used for gas-exchange measurements, following Sakoda et al.(2020). The abaxial epidermis was observed at 250× magnification using an optical microscope. Six images (0.1957 mm² each) were taken per leaf, and stomatal density was calculated using ImageJ software (NIH, Bethesda, MD, United States) by dividing the number of stomata by the area of each image.

#### Proteomic analyses

Total proteins for shotgun proteomics were extracted from 200 mg of rosette powder using the TCA-acetone method described in Langella et al. (2013). The proteomic analyses were realized as described by James et al. (2025).

#### Metabolomic analyses

Metabolic analysis was performed by gas chromatography - mass spectrometry using a Shimadzu GCMS-QP2010SE (single quadrupole) with a Rtx-5MS/integra-Guard capillary column (30 m, 0.25 mm ID, 0.25 µm film thickness / 10 m integrated guard column). Polar metabolites were extracted from 7 mg of lyophilized rosette powder. Sample were derivatized and analyzed by gas chromatography-mass spectrometry as described in Barrit *et al*. (2024). Data were processed using the LabSolutions software (version 4.53, Shimadzu).

#### Nitrate Uptake and nitrogen remobilization to seeds analyses

The rosette uptake experiment was carried out at vegetative and every week at reproductive stage (after the shift to long days), once the inflorescences had reached 2 cm high. For each experiment, six plants per genotype were watered with 10 mM K^15^NO₃ (10 atom% excess) nutritive solution for 48 h and harvested by separating roots, rosettes, and stems (for the reproductive stage).

To quantify nitrogen remobilization to the seeds, plants were labelled at the vegetative stage (one week before the transfer to long-day conditions) providing the 10 mM K¹⁵NO₃ (10 atom% excess) nutritive solution for 24 h. After labeling, sand was thoroughly rinsed with deionized water to remove residual tracer. Plants were then grown until the end of the plant cycle and seed maturity to quantify ^15^N, N and dry weights of each organ separately.

A subsample (0.800–1.200 mg) of each powdered tissue sample was analysed for total nitrogen (N), total carbon (C), and ^15^N abundance using a FLASH 2000 Organic Elemental Analyzer coupled to an isotope ratio mass spectrometer (Thermo Fisher Scientific). Indicators of nitrogen uptake and remobilization efficiencies were calculated following Havé et al. (2017) and Marmagne et al. (2022), and described in our *Arabidopsis Trait Ontology* (Chardon, 2024).

### Statistical analyses

All statistical analyses were performed in R (v4.5.1). Pairwise comparisons between genotypes were conducted using two-tailed Student’s t-tests with wild type as reference. Each experiment was performed twice with at least four biological replicates per genotype per experiment (n ≥ 8 plants per genotype). Data are presented as means ± SD, and significance levels are indicated by stars.

For metabolomic and proteomic datasets, normalized abundance values were centered and scaled (z-score) prior to multivariate analyses. Hierarchical clustering was performed using Ward’s minimum variance method on Euclidean distance matrices, and heatmaps were generated using the ComplexHeatmap package in R. Clusters were defined based on dendrogram structure, and metabolites or proteins were annotated by functional or biochemical categories displayed as row annotations. Relative abundances are shown as color gradients from low to high.

## ACCESSION NUMBERS

Arabidopsis sequence data used in this study are available from TAIR (https://www.arabidopsis.org/): *CLC-A* (AT5G40890), *NRT2.7* (AT5G14570). The barley *HvNRT2.10* sequence (*HORVU.MOREX.r3.7HG0729020*) is available from EnsemblPlants (https://plants.ensembl.org/).

## AUTHOR CONTRIBUTIONS

A.M.., and F.C. conceived the study. A.M., Y.F., B.B., C.C., J.L., and F.C. designed the experimental and analytical approaches and carried out the experiments. A.M., C.M.D. and F.C. performed data analysis. A. M. and F.C. drafted the manuscript. A.M., F.C. and C.M.D. finalized the manuscript. All authors reviewed and approved the final manuscript.

## ACKNOWLEDGEMENTS

We are grateful to Virginie Bréhaut, Patrick Armengaud, and Sophie Filleur for providing seeds of mutant lines. This work has benefited from the support of IJPB’s Plant Observatory platforms PO-Plants.

## SUPPORTING INFORMATION

Supplemental Figure 1: Seed lipid composition.

Supplemental Figure 2: Hierarchical clustering of the leaf metabolome across genotypes.

