## Supplemental Figures S1 and S2 for "Redirecting vacuolar nitrate transport improves nitrogen use efficiency and seed protein content"

### Supplemental information

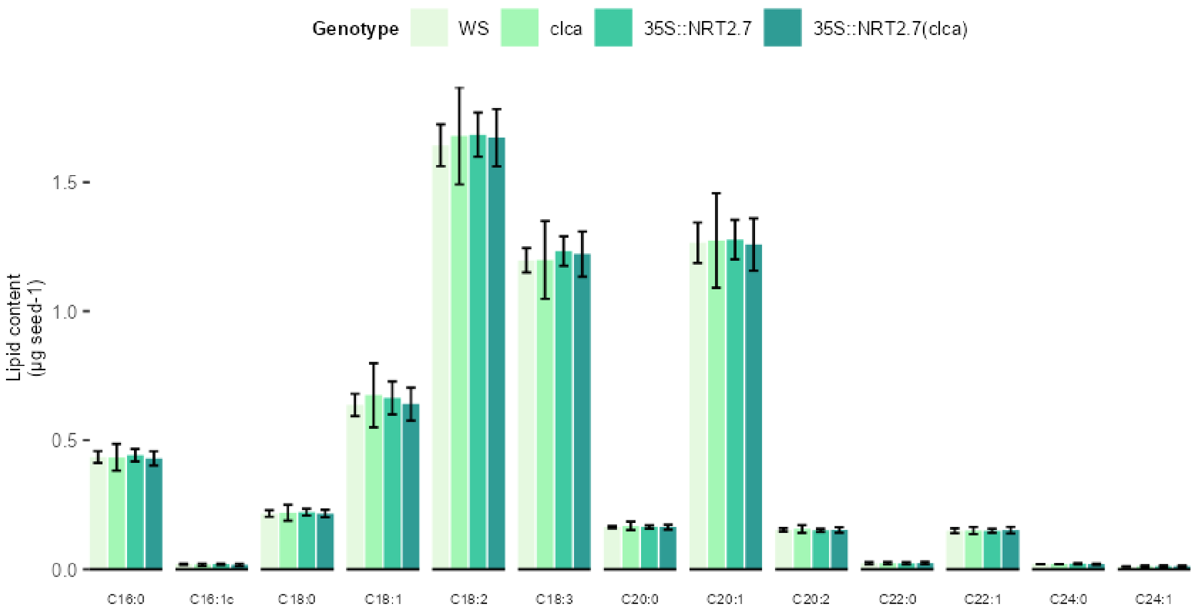


**Supplemental Figure 1: Seed lipid composition**. Fatty acid composition of whole mature seeds from the four ***Arabidopsis thaliana*** genotypes analyzed at maturity. No significant differences in lipid composition were detected among genotypes (Student's *t*-test, *n* = *4*).

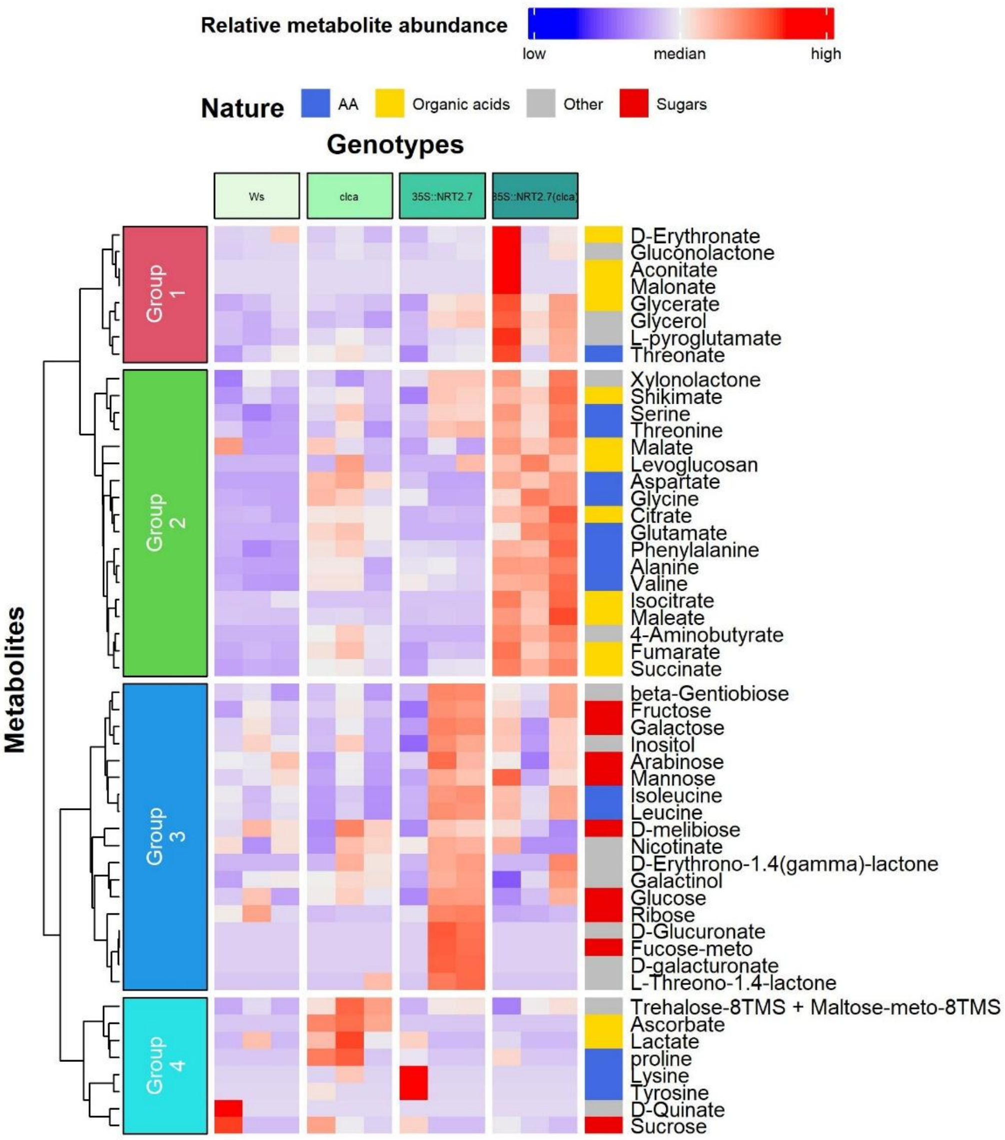


**Supplemental Figure 2: Hierarchical clustering of the leaf metabolome across genotypes.** Heatmap of normalized metabolite abundances measured in vegetative rosettes of WS, *clc-a*, 35S::NRT2.7, and 35S::NRT2.7(*clc-a*) (n = 3 biological replicates per genotype). Metabolites were clustered using Ward.D2 (rows), and grouped into four clusters (Groups 1-4). Metabolite classes (amino acids, sugars, organic acids, others) are indicated by the side annotation.
